# Splice-specific *lyn* knockout mice reveal a dominant function of LynB in preventing autoimmunity

**DOI:** 10.1101/2021.05.03.439514

**Authors:** Ben F. Brian, Monica L. Sauer, Brian Ruis, Jennifer L. Auger, S. Erandika Senevirathne, Whitney L. Swanson, Myra G. Nunez, Branden S. Moriarity, Clifford A. Lowell, Bryce A. Binstadt, Tanya S. Freedman

**Affiliations:** Department of Pharmacology, University of Minnesota; Minneapolis, MN 55455, United States; Graduate Program in Biochemistry, Molecular Biology, and Biophysics; University of Minnesota; Minneapolis, MN, 55455, United States; Center for Genome Engineering, University of Minnesota; Minneapolis, MN, 55455, United States; Department of Pediatrics, Division of Rheumatology, Allergy & Immunology, University of Minnesota; Minneapolis, MN, 55455, United States; Department of Laboratory Medicine, University of California, San Francisco; San Francisco, CA, 94143, United States; Center for Immunology, University of Minnesota; Minneapolis, MN, 55455, United States; Masonic Cancer Center and Center for Autoimmune Diseases Research, University of Minnesota; Minneapolis, MN 55455, United States

## Abstract

The unique roles of the Src-family kinases LynA and LynB in immune activating and inhibitory signaling have eluded definition. Here we report that LynB, the shorter splice product of *lyn*, carries the dominant immunosuppressive function. We used CRISPR/Cas9 gene editing to constrain *lyn* splicing and expression to a single product: LynA^KO^ or LynB^KO^ mice. While activities of both isoforms regulate homeostatic Lyn expression, only LynB protects against autoimmune disease. LynB^KO^ monocytes and dendritic cells are TLR4-hyper-responsive, and TLR4 expression increases with age in LynB^KO^ myeloid and B cells. These changes are accompanied by the development of an inflammatory disease that resembles human systemic lupus erythematosus (SLE). The interplay between LynB and TLR4 likely underlies the autoimmunity risk associated with *LYN* hypomorph and *TLR4* hypermorph alleles.

## Main Text

In myeloid and B cells the Src-family kinase (SFK) Lyn phosphorylates immunoreceptor tyrosine-based activation motifs (ITAMs) and Syk kinase to activate antimicrobial signaling (*1, 2*). Lyn is also inhibitory, dampening ITAM (*2, 3*) and Toll-like receptor (TLR) (*4*) signaling. Hypomorphic polymorphisms of *LYN* are risk alleles for human systemic lupus erythematosus (SLE) (*5, 6*), and patients can have functional deficiencies in Lyn expression, signaling, or trafficking (*7, 8*). Lyn knockout (^KO^) mice develop lupus, with production of anti-nuclear antibodies (ANA), glomerulonephritis, myeloproliferation, and splenomegaly (*9, 10*). Factors controlling whether Lyn initiates pro-inflammatory vs. immunosuppressive signaling have remained a mystery and a barrier to therapeutic development.

Lyn is expressed as the alternatively-spliced isoforms LynA (56 kDa) and LynB (53 kDa) (*11*), which differ only in a 21-residue insert in LynA, encoded by the 3’ region of exon 2 (Fig. 1A, Fig. S1A) (*11*). Using overexpression/reconstitution in Lyn-deficient cells, previous studies have suggested that the roles of LynA and LynB may be non-overlapping (*12, 13*). We have observed that selective degradation of LynA results in signaling loss, and thus also have inferred non-overlapping functions for the two isoforms of Lyn (*14, 15*). To unmask any isoform-specific functions of Lyn, we used CRISPR/Cas9 to generate splice-fixed LynA^KO^ and LynB^KO^ mice.

**Fig. 1.**
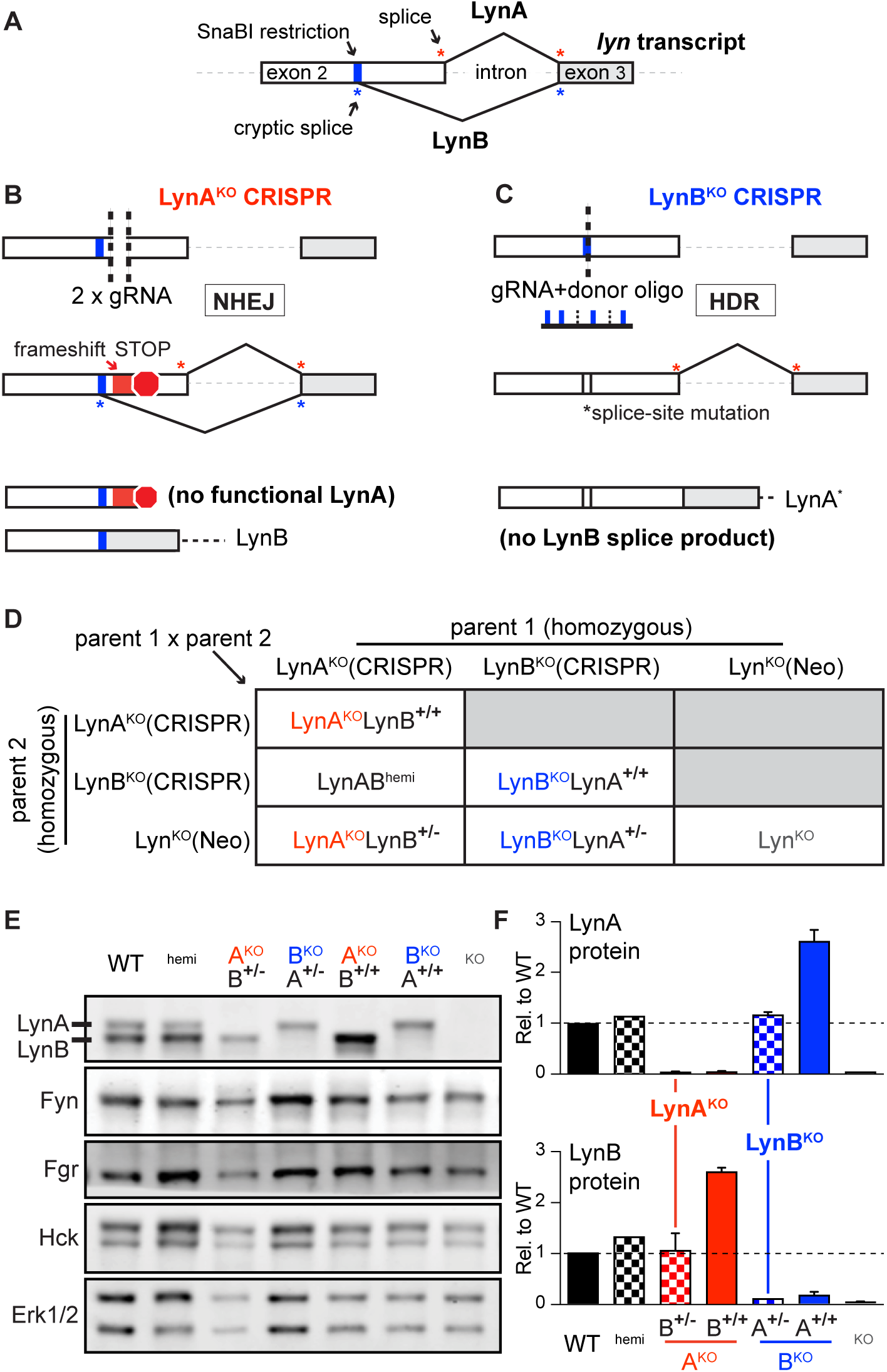
LynA^KO^ and LynB^KO^. **(A)** LynA and LynB splice junctions in WT *lyn*, highlighting a SnaBI site near the cryptic LynB splice donor in exon 2. **(B)** Strategy for LynA^KO^ via double cutting and NHEJ within the LynA insert. **(C)** Strategy for LynB^KO^ via HDR, ablating the LynB splice donor. **(D)** Breeding scheme for LynA^KO^ (A^KO^), LynB^KO^ (B^KO^), LynAB^hemi^ (^hemi^), and total Lyn^KO^ (^KO^) mice, showing parental CRISPR and Neomycin (Neo) (*16*) knockouts. For progeny LynA^KO^LynB^+/+^=biallelic LynB expression, LynA^KO^LynB^+/-^=monoallelic LynB expression (isoforms reversed in LynB^KO^). **(E)** Immunoblot showing SFK expression in WT/knockout BMDMs; Erk1/2 shows loading. **(F)** Quantification (*17*) of LynA and LynB protein; in subsequent figures LynA^KO^LynB^+/-^ and LynB^KO^LynA^+/-^ are referred to as LynA^KO^ and LynB^KO^, respectively.

For LynA^KO^, we injected embryos with Cas9 and two guide (g)RNAs, excising part of *lyn* exon 2 and triggering repair by non-homologous end joining (NHEJ) (Fig. 1B); shortened PCR-screening candidates were sequenced and bred to homozygosity. LynA^KO^(CRISPR) resulted in a frameshift and a premature stop codon in LynA (Fig. S1B), with no effect on LynB (Fig. S1C).

For LynB^KO^, we used a single gRNA and donor oligonucleotide to template homology-directed repair (HDR), inducing one mutation ablating the LynB splice donor and a second (silent) mutation ablating a SnaBI restriction site (Fig. 1C). In LynB^KO^(CRISPR), ablation of the LynB splice donor caused a V24L substitution in LynA (Fig. S2). We performed control experiments to confirm that activation and regulation of LynA^V24L^ was comparable to wild type (WT) LynA (Supplementary Text, Fig. S3A to D). We also generated LynA^KO^LynB^+/+^ x LynB^KO^ LynA^+/+^ F1 (LynAB^hemi^) mice (Fig. 1D, Fig. S3E), which have WT-like expression levels of LynA^V24L^ and LynB (Fig. 1E) and are phenotypically indistinguishable from WT (e.g. Fig. S3F). We therefore consider LynA^V24L^ an acceptable substitute for WT LynA.

LynA and LynB protein were absent as expected in LynA^KO^LynB^+/+^ and LynB^KO^LynA^+/+^ bone-marrow-derived macrophages (BMDMs) (Fig. 1D and E), with preserved interferon (IFN)-γ-dependent upregulation (Fig. S4A and B) (*15*). Expression of the remaining Lyn isoform, however, was elevated 2.5x; other SFKs were unaffected (Fig. 1E and F). This suggests a feedback mechanism that senses and independently tunes basal LynA and LynB expression. To achieve WT-like expression of the remaining isoform, we generated Lyn^KO^ x homozygous LynA^KO^(CRISPR) or LynB^KO^(CRISPR) F1 mice (Fig. 1D), which expressed the remaining Lyn isoform from only one allele, with levels comparable to WT (Fig. 1E and F). These LynA^KO^LynB^+/-^ and LynB^KO^LynA^+/-^ mice are hereafter referred to as LynA^KO^ and LynB^KO^.

Expression of LynA or LynB alone reversed the B-cell deficiency found in total Lyn^KO^ mice (Fig. 2A). Surprisingly, expression of either Lyn isoform also increased CD4^+^ and CD8^+^ T-cell numbers over WT (Fig. 2B); LynAB^hemi^ was indistinguishable from WT. LynB^KO^ spleens had elevated numbers of classical monocytes and trended toward increased patrolling monocytes and granulocytes (Fig. 2C and D). Activation of dendritic cells (DC) drives autoimmune disease in Lyn^KO^ mice (*4, 18–20*), but in contrast to monocytes and granulocytes, DC populations are similar in Lyn^KO^ and WT mice (*19, 21*). LynB^KO^ mice, however, had increased conventional-type-2-DC (cDC2) and plasmacytoid-DC (pDC) numbers (Fig. 2E and F). cDC2 regulate CD4^+^ T-cell activation and promote germinal center formation, while pDC secrete cytokines and drive autoimmune disease (*22*). We assessed spleen organization and found that WT and LynA^KO^ spleens had well-formed lymphoid follicles, whereas LynB^KO^ and Lyn^KO^ had follicular effacement (*23*) (Fig. 2G). Together, these data suggest that LynB has a unique negative-regulatory role, restraining DC and monocyte proliferation and maintaining spleen organization; LynA may therefore have a counterbalancing effect.

**Fig. 2.**
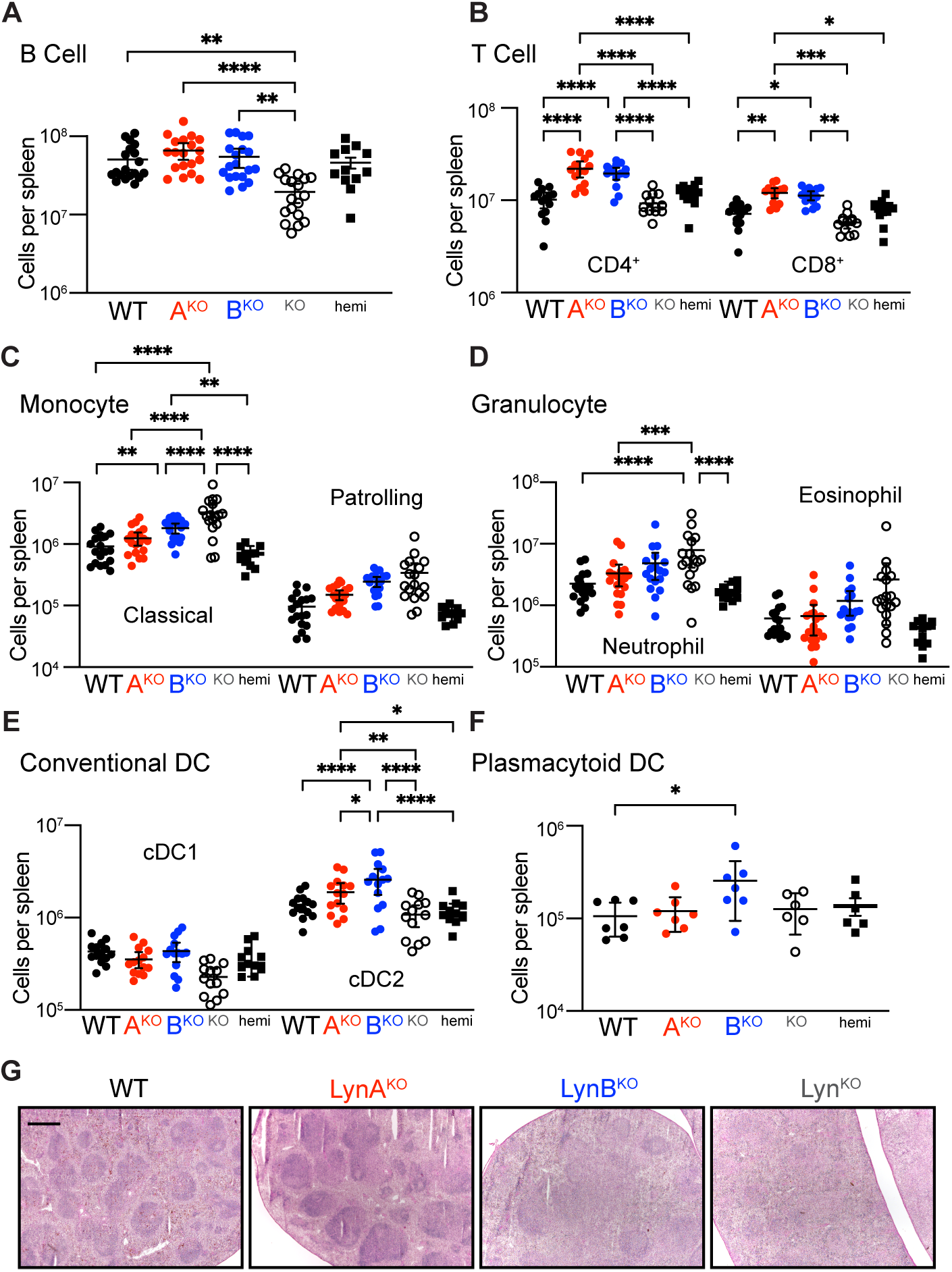
Spleen cellularity and architecture in aged LynB^KO^ mice. Spleens harvested from 8-month-old WT and Lyn knockout mice analyzed via flow cytometry or histology. **(A** to **F)** Cellularity of recovered splenocytes, reported as total number per spleen; points describe data from different individuals compiled from 3 independent cohort analyses. Cell populations include (A) B cell, (B) T cell, (C) monocyte, (D) granulocyte, (E) cDC1, cDC2, and (F) pDC. Statistical tests and cell markers described in the supplement/methods. **(G)** Spleen sections visualized by H&E staining; bar=500 μm.

TLR4 initiates pro-inflammatory signaling in response to lipopolysaccharides (LPS) (*24*). Functional increases in TLR4 signaling, due to increased activity (*25*) or receptor overexpression (*26*), are risk factors for human SLE and drivers of lupus in mice (*25–27*). Lyn can positively or negatively regulate TLR4 signaling in myeloid cells; in DCs it has a primarily negative-regulatory role (*4, 19, 20, 28*). Because DCs are key regulators of the adaptive immune response and drivers of lupus in Lyn^KO^ mice, we investigated TLR4 signaling in bone-marrow-derived dendritic cells (BMDCs) treated with LPS. Like Lyn^KO^ (*4*), LynB^KO^ BMDCs were hyperresponsive to LPS, with increased phosphorylation of IκB kinase (IKK) and Erk1/2 (Fig. 3A); like WT, LynA^KO^ WT BMDCs were less responsive to LPS. LynB, therefore, carries the TLR4-suppressive function historically attributed to Lyn. LynB^KO^ similarly increased LPS-induced production of tumor necrosis factor (TNF) in splenocytes (Fig. 3B). Furthermore, TLR4 expression on circulating B and myeloid cells increased significantly in older Lyn^KO^ and LynB^KO^ mice (Fig. 3C), consistent with observations in human autoimmune diseases (*29*).

**Fig. 3.**
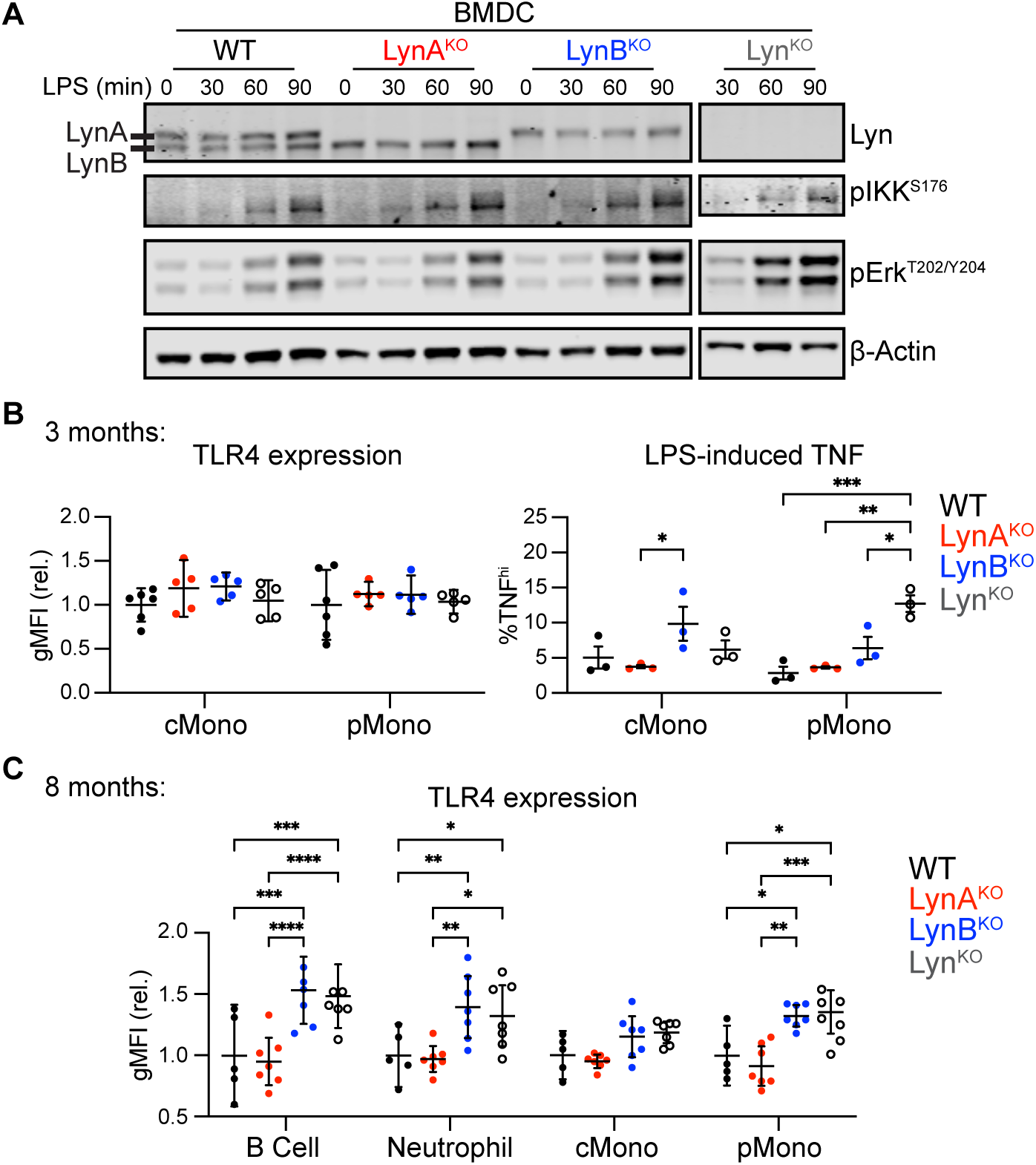
Loss of LynB promotes TLR4 signaling. **(A)** Immunoblots showing phosphorylation of IKK and Erk in response to LPS; β-Actin reflects loading, representative of 3 independent experiments. **(B)** Geometric Mean Fluorescence Intensity (gMFI) showing surface expression of TLR4 in circulating classical and patrolling monocytes in 3-month-old mice. **(C)** Splenocytes from 3-month-old mice treated 4 h with LPS, stained for cell markers/TNF and analyzed via flow cytometry. Points indicate intracellular-TNF^hi^ monocyte populations in preparations from different spleens. **(C)** Blood leukocytes from 8-month-old mice with flow-cytometric quantification of relative TLR4 surface expression.

Like Lyn^KO^ mice (*9, 10, 30*), LynB^KO^ mice lost body mass and developed splenomegaly (Fig. 4A, Fig. S5A and B). LynA^KO^ mice trended toward mild splenomegaly, whereas LynAB^hemi^ mice were indistinguishable from WT (Fig. 4B). Most Lyn^KO^ and LynB^KO^ mice develop severe splenomegaly, in contrast to only 7% of LynA^KO^ mice (Fig. 4C).

**Fig. 4.**
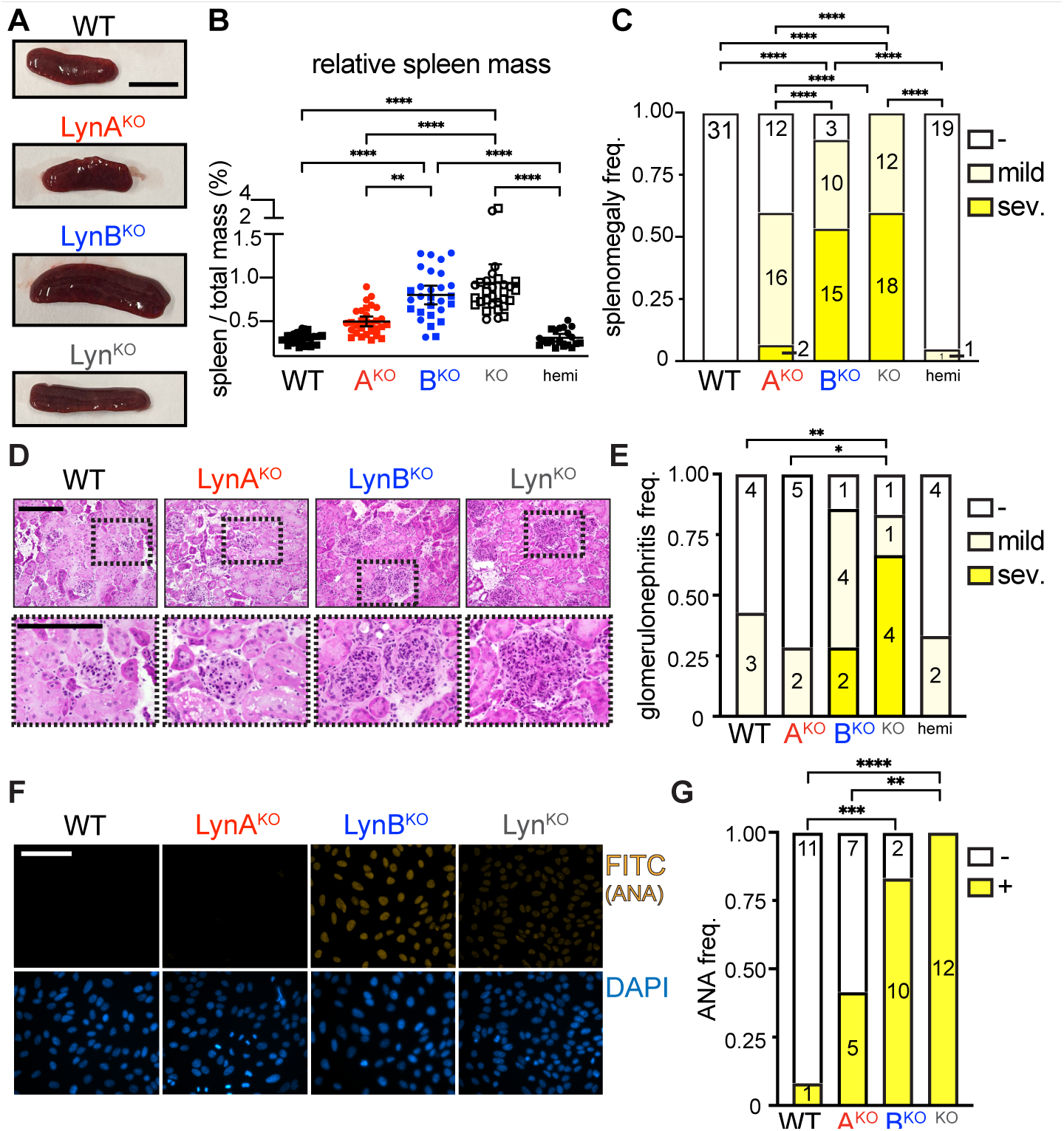
LynB^KO^ is the dominant driver of autoimmune disease. WT and Lyn knockout mice aged 8 months and tested for indicators of autoimmune disease. **(A)** Representative spleens; bar=1 cm. **(B)** Relative spleen mass (spleen mass / total mass) of male (square) and female (circle) mice. **(C)** Frequency of mice with no (-, spleen mass <0.45% total), mild (0.45-0.75%), or severe (sev.) splenomegaly (>0.75%). **(D)** H&E-stained kid-ney sections scored for glomerulonephritis by a blinded pathologist; bars=200 μm, bottom enlarged from boxes. **(E)** Frequency of glomerulonephritis; numbers are scores from individual mice. **(F)** Serum probed for IgG-ANA; bar=100 μm, representative of 3 experiments. **(G)** Frequency of ANA negativity (-) and positivity (+); numbers are sera from individual mice.

Lyn^KO^ and LynB^KO^ kidney sections showed severe immune infiltration, expanded glomeruli, and tubular inflammation (Fig.4D and E), in contrast to no/mild inflammation in WT, LynA^KO^ and LynAB^hemi^ mice, and most Lyn-B^KO^ and Lyn^KO^ mice produced ANA, unlike LynA^KO^ and WT (Fig. 4F and G). No sex-specific differences were observed (Fig. S6). Together, these data indicate that the lupus-like disease in aging Lyn^KO^ mice is caused by the loss of LynB function, suggesting that LynB is the dominant immunosuppressive/protective isoform.

Although it has long been appreciated that Lyn is a critical regulator of immune signaling, dissecting the contributions of the two isoforms, LynA and LynB, has been stymied by the lack of experimental tools. LynB has no unique sequence, making development of LynB-specific antibodies and RNA silencing reagents infeasible; LynB production from a cryptic, intra-exon splice site rendered traditional knockout and recombination/excision approaches intractable due to residual effects on LynA.

We pioneered two CRISPR/Cas9 strategies to create splice-fixed LynA- or LynB-only mouse strains: (i) deleting/frameshifting the unique LynA insert (LynA^KO^) or (ii) ablating the LynB splice donor (LynB^KO^), both of which can express the remaining isoform from one allele or two, for physiological or supraphysiological expression. We found that protein levels of LynA and LynB are coregulated by a feedback mechanism that senses the basal activity of either isoform. This Lyn-specific feedback, independent of other SFKs, highlights the unique importance of balancing the activating and inhibitory functions of Lyn for proper control of cellular responses and maintenance of immune homeostasis. While expression of LynA or LynB alone is sufficient to prevent the B-cell lymphopenia observed in Lyn^KO^ mice, LynB supplies the primary immunoregulatory activity that protects against autoimmune disease, with its loss leading to increased TLR4 signaling and both innate- and adaptive-immune dysregulation. Disease indicators and hyperresponsive signaling in LynB^KO^ mice and cells are comparable to total Lyn^KO^, suggesting that LynA and LynB functions are distinct, not additive. This supports a model in which LynA tunes the sensitivity of cells to pro-inflammatory activation (*14, 15*) and LynB directs suppressive signaling. Definition of LynA-specific and LynB-specific signaling pathways and their respective contributions to immune regulation will inspire new ways of understanding and manipulating these signaling pathways to restore immune balance in SLE and other autoimmune patients.

## Materials and Methods

### Citation Practices

Please refer to our previous publications (*30, 14, 15*) for comprehensive referencing of materials, methodologies, and primary literature.

### Generation of LynA^KO^ and LynB^KO^ mice

LynA^KO^ was induced using two gRNAs to delete 77 bp encompassing portions of the unique LynA insert in mouse lyn exon 2 (5’-GAUCUCUCACAUAAAUAGUU-3’) and the following intron 2 (5’-CCAUGCUCCGAUCCUACUGU-3’). NHEJ then induced a frame shift that resulted in a premature stop codon after amino-acid residue 77.

LynB^KO^ was induced using one gRNA (5’-GUUCGGUCAGUAUUACGUAC-3’) to make a cut near the LynB splice site in exon 2. A donor oligonucleotide (5’-AAAAGGAAAGACAATCTCAATGACGATGAAGTAGAT-TCGAAGACTCAACCAcTgCGTAA-TACTGACCCAACTATTTATGTGAGAGATCCAACGTCCAATAAACAGCAAAGGCCAGTAAG-3’) was supplied to induce two single-nucleotide substitutions via HDR (lower-case letters in the sequence) that ablated the LynB splice site and a SnaBI restriction site. gRNA’s were designed using CRISPOR.org.

3-week-old C57BL/6J female mice were purchased from The Jackson Laboratory and adapted to University of Minnesota animal facilities 3-4 days before the first hormone injection. The stud C57BL/6J male mice were obtained from the Jackson Laboratory directly. The recipient females CD-1 mice at 38-49 days old was purchased from the Charles River Laboratory. The animal protocols used for all mutant mice generation were approved by the Institutional Animal Care and Use Committee (IACUC).

Microinjection of embryos: The C57BL/6J females at 3-4 weeks old were superovulated by intraperitoneal injection of 5 IU pregnant mare serum gonadotropin, followed by injection of 5 IU human chorionic gonadotropin 48 hrs. later (both hormones from National Hormone & Peptide Program, Torrance, CA, USA), and they were immediately crossed to C57BL/6J stud males. The next day, mouse embryos were obtained from superovulated C57BL/6J females. For LynA^KO^, the embryos were injected with mixture of 30 ng/ul Cas9 protein, 3.5 ng/ul sgRNA each of 2 gRNAs. For LynB^KO^, the embryos were injected with mixture of 30 ng/ul Cas9 protein, 3.5 ng/ul of gRNA, and 7 ng/ul of single-stranded oligonucleotide (120bp) donor as the point mutation template. CRISPR reagents were designed by University of Minnesota Genome Engineering Shared Resources and purchased from IDT. All CRISPR reagents were tested and validated on NIH 3T3 cells before the embryo injection. And all the CRISPR reagents were resuspended in the injection buffer (10 mM Tris-HCl, 0.25 mM EDTA, pH 7.4) for the embryo injection

Toe DNA from the resulting pups was isolated using DNeasy Blood & Tissue Kit (Qiagen, Hilden, Germany) and subjected to and intermediate topo cloning step (Invitrogen, Carlsbad, CA) to insert PCR products into a plasmid for PCR sequencing (GeneWiz, South Plainfield, NJ) and digestion. A 500 bp segment in wild-type *lyn* encompassing the LynB splice site and the LynA unique insert was amplified using the following PCR primers: Forward 5’-acaaccgagatgtcctgct-3’ Reverse 5’-agccagattatccctaaaatctctaca-3’. SnaBI (New England Biolabs, Ipswich, MA) cleavage of this product (recognition site TAC/GTA) generated two 250 bp fragments. In LynA^KO^ a Cas9 double-cut deletion yields a shorter (~423 bp) PCR product that retains sensitivity to SnaBI cleavage. In LynB^KO^ an HDR-induced mutation in the SnaBI site yields a 500 bp PCR product insensitive to SnaBI.

### Other mouse strains and housing

C57BL/6-derived Csk^AS^ mice are hemizygous for the Csk^AS^ BAC transgene on a Csk^-/-^ background, as described previously (*15*). Lyn^KO^ mice (*29*) were maintained on a C57BL/6 background. All mice were housed in specific pathogen-free conditions and genotyped using real-time PCR (Transnetyx, Inc, Memphis, TN). All animal use complies with University of Minnesota (UMN) and National Institutes of Health (NIH) policy (Animal Welfare Assurance Number A3456-01). UMN is accredited by AAALAC, experiments involving mice were approved by the UMN Institutional Animal Care and Use Committee (IACUC, protocol # 1603-33559A). Animals are kept under supervision of a licensed Doctor of Veterinary Medicine and supporting veterinary staff under strict NIH guidelines.

### Jurkat cell lines and transfection

The Jurkat T-cell strain JCaM1.6 (Lck-deficient) was a gift from the laboratory of Y. Shimizu (University of Minnesota). Cell lines were authenticated by STR profiling and tested negative for mycoplasma (ATCC, Manassas, Virginia). JCaM1.6 was cultured in RPMI-1640 medium supplemented with 5-10% fetal bovine serum (FBS) (Omega Scientific, Inc, Tarzana, CA) and 2 mM glutamine, penicillin and streptomycin (Sigma-Aldrich, St. Louis, MO). JCaM1.6 cells were transiently transfected via electroporation. Briefly, cells were grown overnight in antibiotic-free RPMI-1640 medium supplemented with 10% FBS (Omega Scientific) and 2 mM glutamine (RPMI10). Batches of 15 M cells were resuspended in RPMI10 with 10–15 μg plasmid DNA per construct. Cells were rested, electroporated at 285 V for 10 ms in a BTX square-wave electroporator (Harvard Apparatus, Holliston, MA), resuspended in RPMI10, and allowed to recover overnight. One million live cells were then resuspended in phosphate-buffered saline (PBS), rested for 30 min at 37°C, and stimulated.

### Preparation of BMDMs

Bone marrow was extracted from femura/tibiae of mice. After hypotonic lysis of erythrocytes, BMDMs were derived on untreated plastic plates (BD Falcon, Bedford, MA) by culturing in Dulbecco’s Modified Eagle Medium (Corning Cellgro, Manassas, VA) containing approximately 10% heat-inactivated FBS (Omega Scientific, Tarzana, CA), 0.11 mg/ml sodium pyruvate (UCSF Cell Culture Facility), 2 mM penicillin/streptomycin/L-glutamine (Sigma-Aldrich, St. Louis, MO), and 10% CMG-12-14-cell-conditioned medium as a source of M-CSF. After 6 or 7 days, cells were resuspended in enzyme-free ethylenediaminetetraacetic acid (EDTA) buffer and replated in untreated 6-well plates (BD Falcon) at 1 M cells per well in unconditioned medium ±25 U/ml IFN-γ (Peprotech, Rocky Hill, NJ) (*14, 30*).

### Flow cytometry

Spleens were excised from mice and cut into <1mm pieces before being digested for 30 min in RPMI 2% FBS, 0.2 mg/mL collagenase type IV (Sigma-Aldrich, St. Louis, MO), 37.5 ug/mL DNAse I (Worthington Biochemical, Lakewood, NJ), 1 U/mL Heparin D Sodium (Sigma-Aldrich, St. Louis, MO), 1 mM CaCl_2_ and 1 mM MgCl_2_. Single-cell suspensions were prepared by filtering digested spleens through 70 μm filters (Celltreat, Pepperell, MA). Antibody master mixes were prepared in PBS 2% FBS, 2mM EDTA. Antibodies used are enumerated in Table 1. For myeloid-cell-staining, cells were stained in 100 μL master mix at 37 °C for 30 min. For T and B cell panels, cells were stained at 4 °C for 30 min. Following intracellular staining, cells were washed with permeabilization buffer and resuspended in PBS 0.5% paraformaldehyde. Data was acquired using a BD Fortessa X-30 and analyzed using FlowJo software.

## Antibodies used in flow cytometry

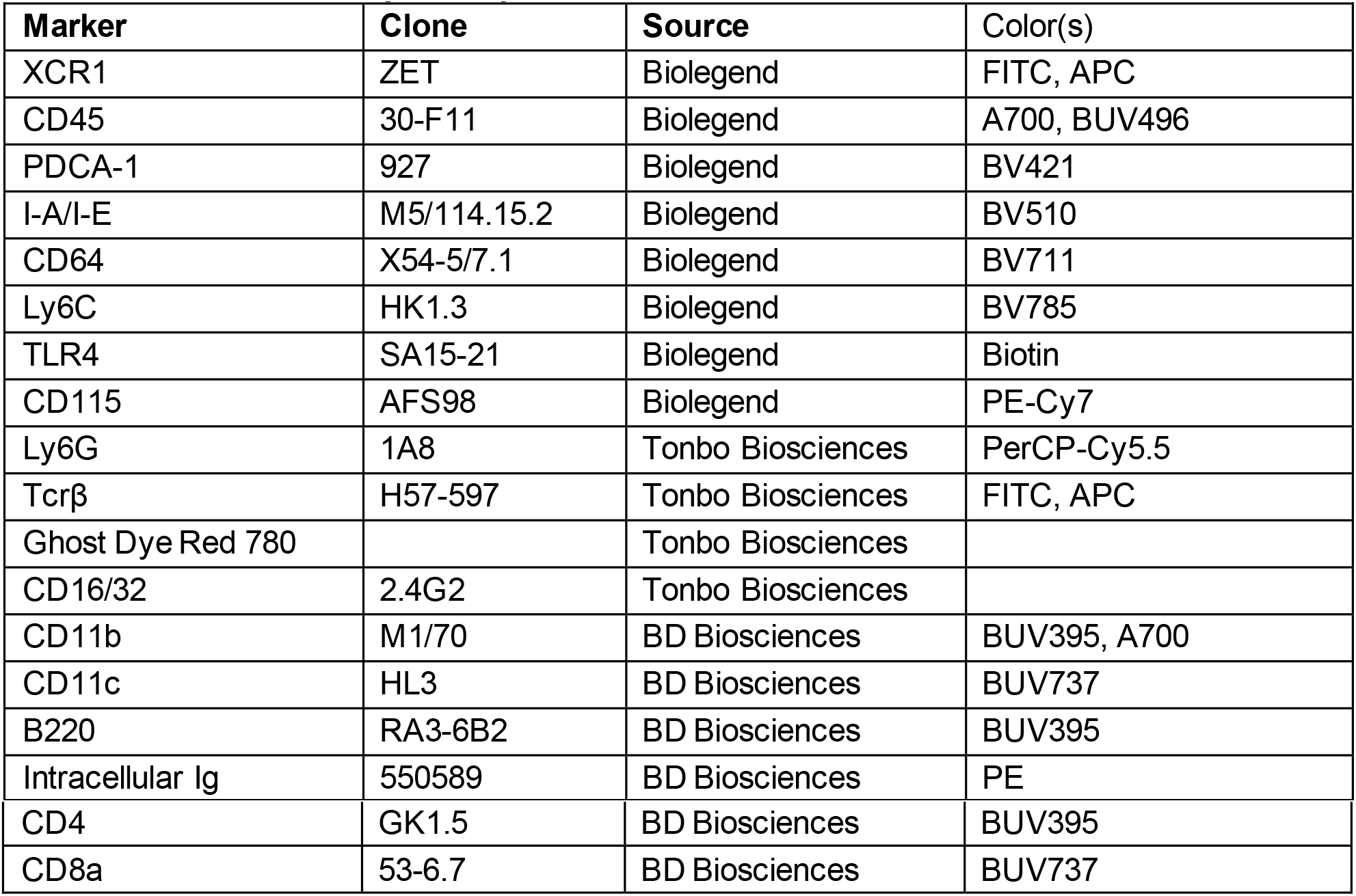

| **Marker** | **Clone** | **Source** | **Color(s)** |
| --- | --- | --- | --- |
| XCR1 | ZET | Biolegend | FITC, APC |
| CD45 | 30-F11 | Biolegend | A700, BUV496 |
| PDCA-1 | 927 | Biolegend | BV421 |
| I-A/I-E | M5/114.15.2 | Biolegend | BV510 |
| CD64 | X54-5/7.1 | Biolegend | BV711 |
| Ly6C | HK1.3 | Biolegend | BV785 |
| TLR4 | SA15-21 | Biolegend | Biotin |
| CD115 | AFS98 | Biolegend | PE-Cy7 |
| Ly6G | 1A8 | Tonbo Biosciences | PerCP-Cy5.5 |
| Tcrβ | H57-597 | Tonbo Biosciences | FITC, APC |
| Ghost Dye Red 780 |  | Tonbo Biosciences |  |
| CD16/32 | 2.4G2 | Tonbo Biosciences |  |
| CD11b | M1/70 | BD Biosciences | BUV395, A700 |
| CD11c | HL3 | BD Biosciences | BUV737 |
| B220 | RA3-6B2 | BD Biosciences | BUV395 |
| Intracellular Ig | 550589 | BD Biosciences | PE |
| CD4 | GK1.5 | BD Biosciences | BUV395 |
| CD8a | 53-6.7 | BD Biosciences | BUV737 |

### Histology

Kidneys were snap-frozen in optimal cutting temperature compound and 5-μm sections were H&E stained. The presence and severity of nephritis was evaluated in a blinded fashion (as previously described by one of the authors. Scoring of glomerulonephritis and interstitial nephritis on a 0-3 scale (0=absent, 1=mild, 2=moderate, 3=severe) for glomeruli was based on glomerular size, glomerular hypercellularity, and presence of glomerular sclerosis, and for interstitial disease was based on the degree of inflammatory infiltrate and alteration in tissue architecture.

### Anti-nuclear antibody staining

Kallestad HEp-2 (Bio-Rad, Cat. No. 30472) was used to detect Anti-Nuclear antibody staining according to the manufacturer’s instruction, modified to detect mouse IgG. Briefly, wells were incubated with 40 μL mouse serum, diluted 40x in phosphate-buffered saline + 1% Bovine-serum albumin, at room temperature for 20 min. After incubation, slides were washed in PBS for 10 minutes and incubated with 30 μL FITC-labeled goat-anti-mouse IgG (Jackson ImmunoResearch, Cat. No. 115-095-164) diluted 200x and DAPI for 20 min. at room temperature, followed by a 10 min. PBS wash. Slides were then mounted and coverslipped and viewed on an Olympus BX51 fluorescent microscope equipped with a digital camera and DP-BSW software (Olympus). Images were quantified, pseudocolored, and merged using ImageJ software (NIH) with the Fiji plugin.

### Splenocyte stimulation ex vivo

Spleens were excised from mice and placed in ice-cold RPMI, 2% FBS, 2 mM penicillin and streptomycin. Spleen tissue was mechanically disrupted using a 100-μm cell strainer (Corning, Corning, NY) on a 50 ml tube, using a plunger of a 1 ml syringe. During and following processing of the spleen, strainers were flushed with ice-cold RPMI to acquire all cells. After processing, single-cell suspensions were counted and kept at 4°C before proceeding with the in vitro stimulation. Stimulation and staining of cells were performed in 96-wells V-bottom plates (Nunc, Roskilde Denmark), using 2×10^6^ cells per well in 50 μL RPMI. Cells were placed in a 5% CO_2_ 37 °C incubator for 1 hr prior to treatment. LPS (Invivogen, San Diego, CA) was diluted in RPMI 2-5% FBS and added by pipetting 50 μl (at 2x concentration) to the wells containing 50 μl of sample. After 30 min, Brefeldin A and monensin (eBioscience, San Diego, CA) were added. For the last 30 min of stimulation, Live/Dead stain (Tonbo Biosciences, San Diego, CA) was added. Signaling was quenched at 4 h by adding paraformaldehyde (PFA) to a final concentration of 1.6% and fixing at room temperature for 15 min in the dark. Following fixation, cells were washed twice with BD Biosciences Fix/Perm diluent and by centrifuging at 0.4 x g for 5 min. and dumping supernatant. Cells were then mixed with staining master mix diluted in BD Biosciences Fix/Perm diluent overnight at 4 °C. Following intracellular staining, cells were washed twice with permeabilization buffer and resuspended in PBS, 0.5% PFA. Data was acquired using a BD Fortessa X-30 and analyzed using Flowjo.

### Preparation of BMDCs

Bone marrow was extracted from femura/tibiae of mice. After hypotonic lysis of erythrocytes, BMDCs were derived on 10 cm tissue-culture-treated, plastic plates (Celltreat, Pepperell, MA) in Dulbecco’s Modified Eagle Medium (DMEM) (Corning Cellgro, Manassas, VA) with ~10% heat-inactivated FBS (Omega), 0.11 mg/ml sodium pyruvate (UCSF Cell Culture Facility), 5 mM penicillin/streptomycin/L-glutamine (Sigma-Aldrich, St. Louis, MO), 10 ng/mL murine GM-CSF (PeproTech, Cranbury, NJ), and 10 ng/mL murine IL-4 (PeproTech, Cranbury, NJ). Media was replaced on day 3 or 4. After 6 or 7 days, cells were resuspended in enzyme-free ethylenediaminetetraacetic acid (EDTA) buffer and replated in TC-treated 12-well plates (Celltreat, Pepperell, MA) at 1 M cells per well in 10%FBS-DMEM.

### BMDC stimulation

BMDCs were replated in 12-well tissue-culture-treated plates and rested overnight in a 37°C incubator. LPS (Invivogen) was added to BMDCs at a final concentration of 100 ng/ml and incubated at 37°C for 30, 60, or 90 min. Samples were collected and prepared for immunoblotting using a standard protocol (*30*).

### Immunblotting

0.025 M cell equivalents were run in each lane of a 7% Nupage Tris-Acetate gel (Invitrogen, Carlsbad, CA) and then transferred to an Immobilon-FL PVDF membrane (EMD Millipore, Burlington, MA). REVERT Total Protein Stain (TPS, LI-COR Biosciences, Lincoln, NE) was used according to the standard protocol to quantify lane loading. After destaining, membranes were treated with Odyssey Blocking Buffer (TBS) for 1 hr. Blotting was performed using standard procedures, and blots were imaged on an Odyssey CLx near-infrared imager (LI-COR). Primary Antibodies: Mouse anti-β-Actin (8H10D10) (Cell Signaling, Cat. #3700, 1:5000), Rabbit anti-pErk1/2 (phosphoThr202/phosphoTyr204, D13.14.4E) (Cell Signaling, Cat. #4370, 1:5000), Rabbit anti-p-IKKα/β (Ser176/180, 16A6) (Cell Signaling, Cat. #2697, 1:1000), Mouse anti-LynA + LynB (Lyn-01) (Abcam, Cat. #ab1890, 1:1000). Secondary Antibodies were either IRDye (680/800) Donkey anti-Mouse IgG or Donkey anti-Rabbit IgG (Li-COR, Biosciences, 1:10000).

### Cell Markers in Fig. 2

(A) B cell (B220^+^), (B) CD4^+^ and CD8^+^ T cell (B220-NK1.1-TCRβ^+^), (C) classical monocyte (Ly6G^-^CD64^+^MerTK^-^ Ly6C^+^), patrolling monocyte (Ly6G^-^CD64^+^MerTK^-^Ly6C^-^), (D) neutrophil (Ly6G^high^), eosinophil (CD64^-^CD11c^-^ SSC^hi^), (E) cDC1 (Ly6G^-^CD64^-^CD11c^+^MHCII^hi^XCR1^+^CD11b^b^), cDC2 (Ly6G^-^CD64^-^ CD11c^+^M HCII^hi^XCR1^lo^CD11b^hi^), and (F) pDC (Ly6G^-^CD64^-^CD11c^+^PDCA-1^+^Ly6C^+^).

### Statistical Analyses

Fig. 2B- Significance (Sig.) assessed using 1-way ANOVA with Tukey’s multiple comparison test (ANOVA_1w_-Tukey); error bars reflect 95% confidence intervals (95%CI), ****P<0.0001, ***P<0.0002, **P<0.0021, *P<0.0332.

Fig. 2C- Sig. for raw contingency data assessed via two-sided Fisher’s exact test (Fisher_2side_) from binarized pools of no/mild vs. severe splenomegaly without correction for multiple comparisons; with Bonferroni correction significance is preserved except for [LynB^KO^ vs. F1] and [Lyn^KO^ vs. F1].

Fig. 2E- Sig. for raw contingency data assessed via Fisher_2side_ from binarized pools of no/mild vs. severe glomerulonephritis without correction for multiple comparisons; with Bonferroni correction significance is preserved except for [LynA^KO^ vs. Lyn^KO^].

Fig. 2G- Sig. for raw contingency data assessed via Fisher_2side_ from - vs. + ANA without correction for multiple comparisons; with Bonferroni correction, significance is preserved except for [LynA^KO^ vs. Lyn^KO^].

Fig. 3A to F- Sig. from ANOVA_1w_- or ANOVA_2w_-Tukey): ****P<0.0001, ***P<0.0002, **P<0.0021, *P<0.0332; error = 95%CI.

Fig. 4B and C- Sig. from ANOVA_1w_- or ANOVA_2w_-Tukey): ****P<0.0001, ***P<0.0002, **P<0.0021, *P<0.0332.

## Acknowledgments

We thank Anindya Bagchi, Yun You, University of Minnesota Mouse Genetics Laboratory and University of Minnesota Genome Engineering Shared Resource for technical assistance. Many thanks also to Kaylee Schwertfeger, Michael Farrar, Carol Lange, John Connett and Marc Jenkins for valuable feed-back and discussion of the project or manuscript.

## Author contributions

Conceptualization: BFB, TSF

Methodology: BFB, JLA, BLR, BSM, CAL, BAB, TSF

Investigation: BFB, MLS. JLA, MGN, SES, WLS, TSF

Visualization: BFB, JLA, TSF

Funding acquisition: BFB, TSF

Project administration: TSF

Supervision: TSF

Writing – original draft: BFB, TSF

Writing – review & editing: BFB, MLS, JLA, MGN, SES, WLS, BLR, BSM, CAL, BAB, TSF

## Data and materials availability

All data are available in the main text or the supplementary materials.

## Supplementary Text

To confirm that V24L substitution does not impair LynA, wild-type (WT) LynA- and LynA^V24L^-initiated signaling was tested in JCaM1.6 (Jurkat T) cells, which depend on introduced LynA for signal initiation (*14*). Specific inhibition of cotransfected memCsk^AS^ with the small molecule 3-IB-PP1 induces spontaneous SFK/LynA activation and downstream signaling (*14*). LynA^V24L^ activation led to Erk phosphorylation (pErk) similarly to WT (Fig. S3A and B). To confirm that V24L substitution does not impair activation-induced LynA degradation, we induced SFK/LynA activation by treating bone-marrow-derived macrophages (BMDMs) from transgenic Csk^AS^ (*14*) Csk^AS^LynB^KO^ mice with 3-IB-PP1. Rates of degradation of WT Lyn and LynA^V24L^ were indistinguishable, demonstrating that the V24L substitution does not impair Y32 phosphorylation, interaction with c-Cbl, or other signal-initiation and regulatory machinery (Fig. S3C and D).

---

**Fig. S1.**
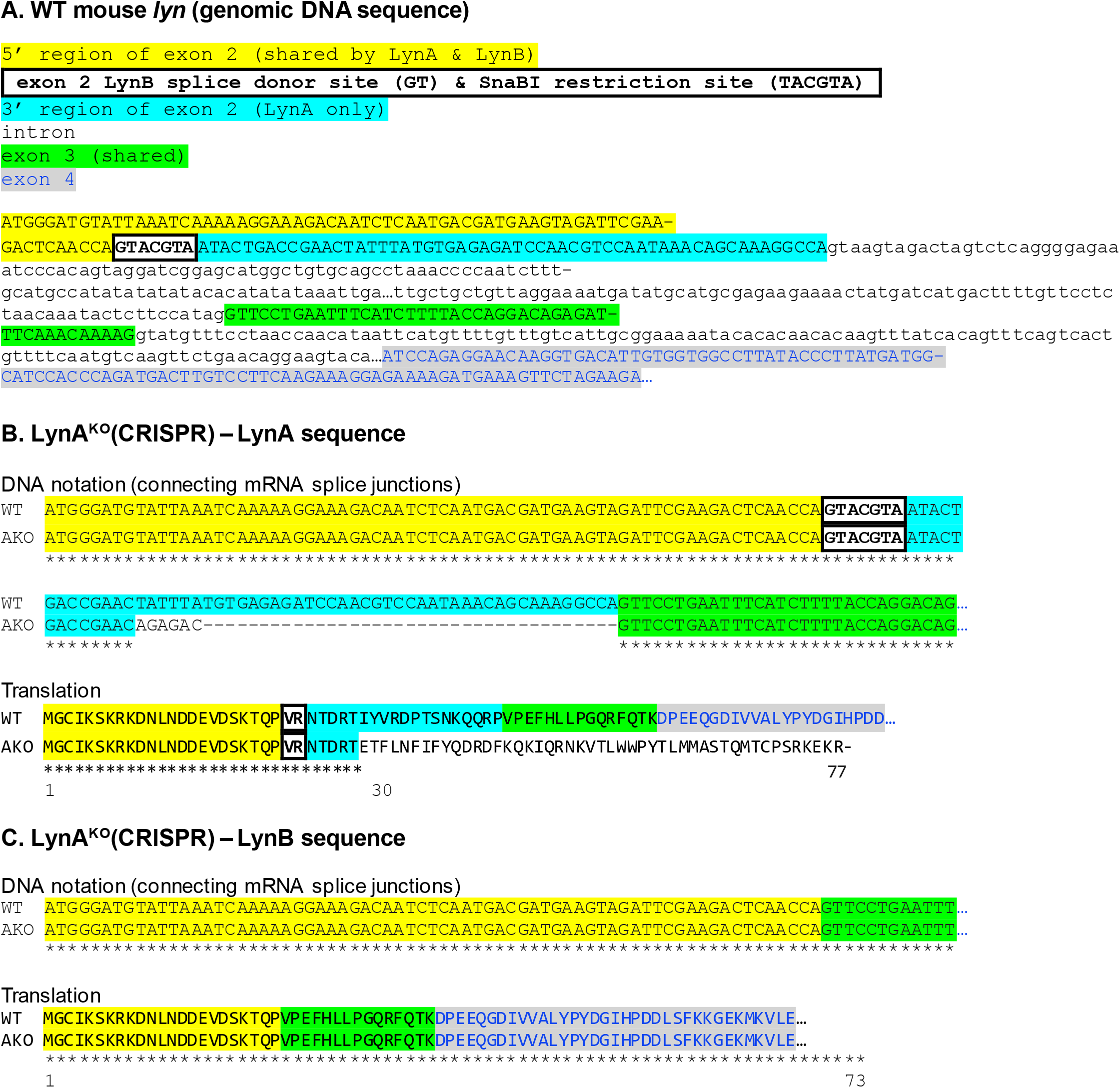
Nucleotide and protein sequence generated by LynA^KO^(CRISPR). **(A)** Genomic sequence of exon 2 (yellow) and exon 3 (green) of wild-type (WT) mouse *lyn*, illustrating the cryptic LynB splice site (boxed) and LynA unique region (cyan). **(B)** LynA^KO^ (AKO) sequences following CRISPR/Cas9 gene editing, illustrating the frameshift and premature stop codon. Nucleotide sequences are shown in DNA notation for ease of comparison to genomic sequence above. Translated sequences were generated using Expasy Translate Tool (https://web.expasy.org/translate/). **(C)** The sequence of LynB is unaffected by these mutations.

**Fig. S2.**
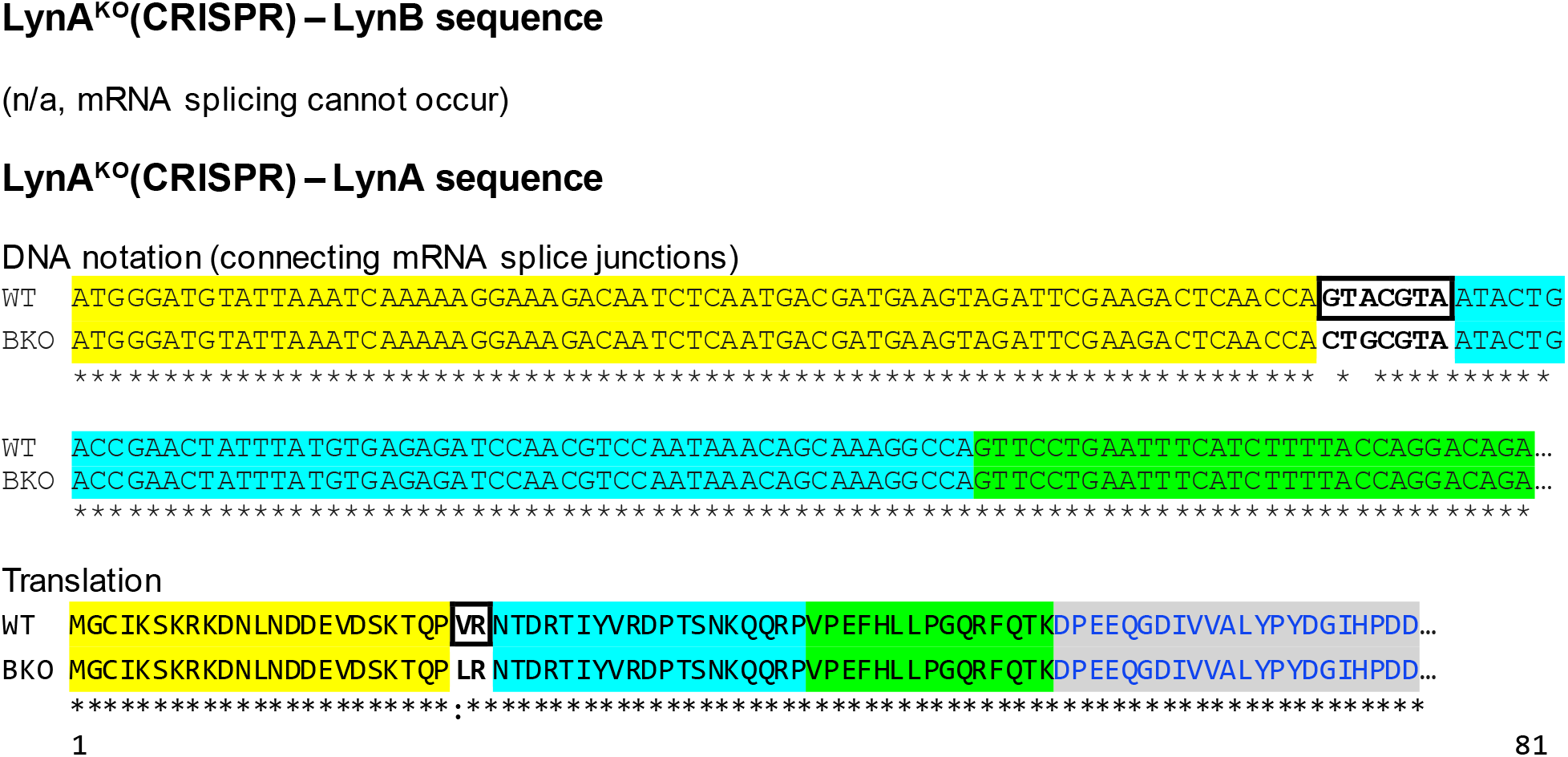
Nucleotide and protein sequence generated by LynB^KO^(CRISPR). LynB is not expressed in the LynB^KO^ due to a splice-site mutation. LynA has a single amino-acid substitution (V24L).

**Fig. S3.**
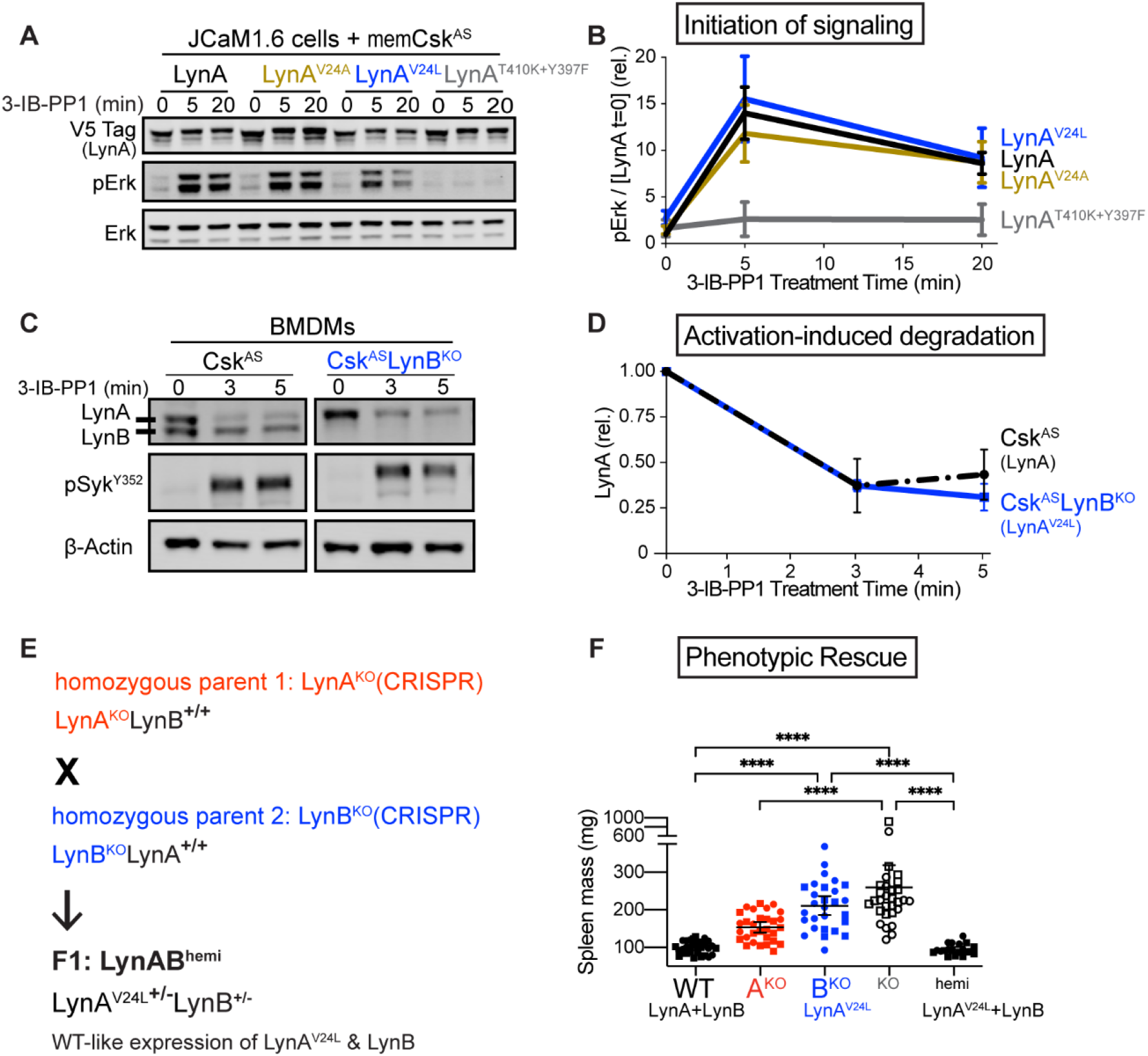
Function is preserved in LynA^V24L^. **(A)** Representative immunoblot of lysates from JCaM1.6 cells transiently transfected with cDNAs encoding LynA and memCsk^AS^ and treated with 3-IB-PP1. Phosphorylation of Erk (pT202/pY204) occurs downstream of LynA (WT and V24L)-initiated signaling, in contrast to kinase-dead LynA^T410K+Y397F^; total Erk reflects protein loading. **(B)** Quantification of Lyn-dependent Erk phosphorylation during 3-IB-PP1 treatment, corrected for the initial amount of transfected LynA protein and shown relative to WT basal. **(C)** Representative immunoblot of lysates from LynA-expressing Csk^AS^ and LynA^V24L^-expressing Csk^AS^LynB^KO^ BMDMs treated with 3-IB-PP1. Syk Y352 is an SFK substrate, and β-Actin reflects protein loading. **(D)** Quantification of LynA degradation in Csk^AS^ and Csk^AS^LynB^KO^ BMDMs. **(E)** Lyn protein expression in parental and LynAB^hemi^ mice. **(F)** Protection of LynA^V24L^-expressing LynAB^hemi^ from the mild splenomegaly occurring in LynA^KO^ mice at 8 months.

**Figure S4.**
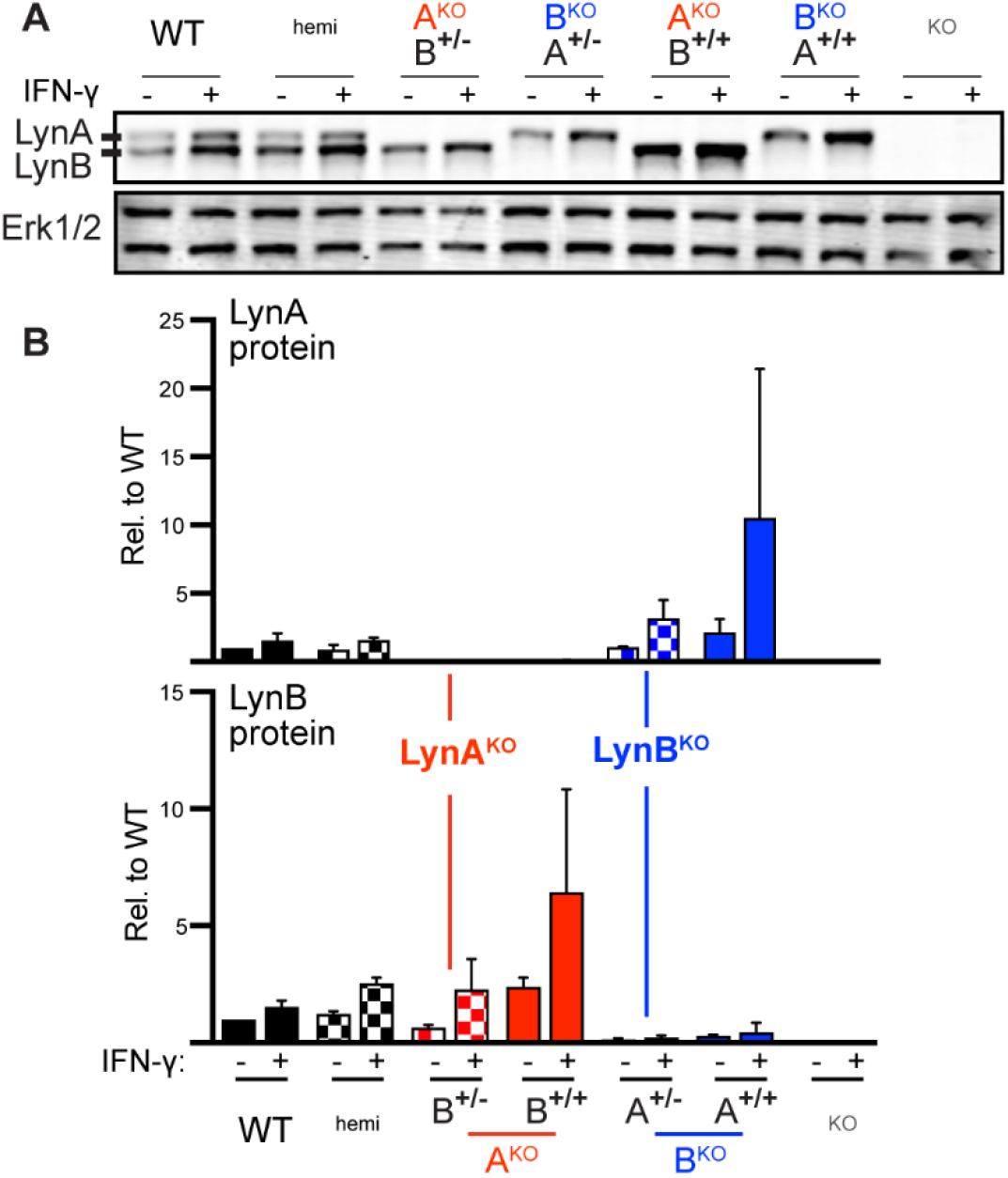
Lyn upregulation in response to IFN-γ is preserved in LynA^KO^ and LynB^KO^ BMDMs. **(A)** Representative immunoblot of WT and Lyn knockout BMDMs rested (-) or incubated overnight with low-dose IFN-γ (+). **(B)** Quantification of LynA and LynB protein in BMDMs before and after IFN-γ treatment; n=2.

**Figure S5.**
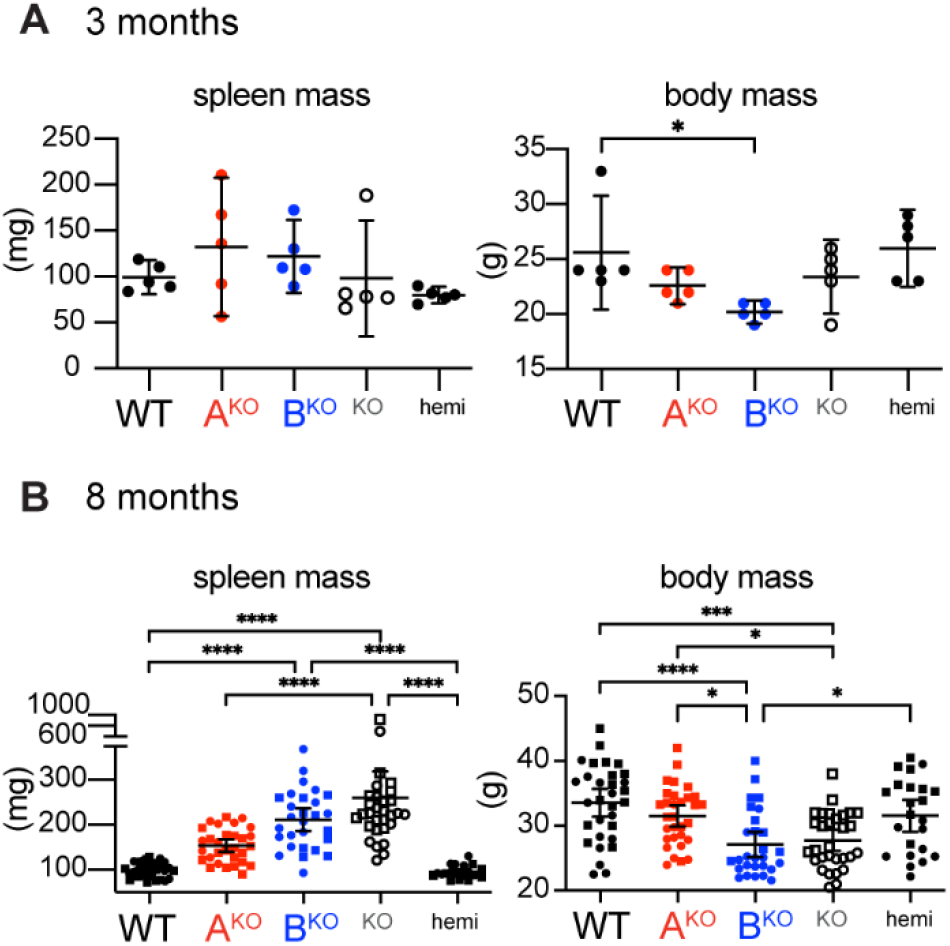
Splenomegaly and weight loss develop over time in LynB^KO^ mice. Spleen and body mass in mice aged **(A)** 3 months or **(B)** 8 months. Sig. ANOVA_1_-Tukey: ****P<0.0001, ***P<0.0002, **P<0.0021, *P<0.0332. Error bars: 95%CI.

**Figure S6.**
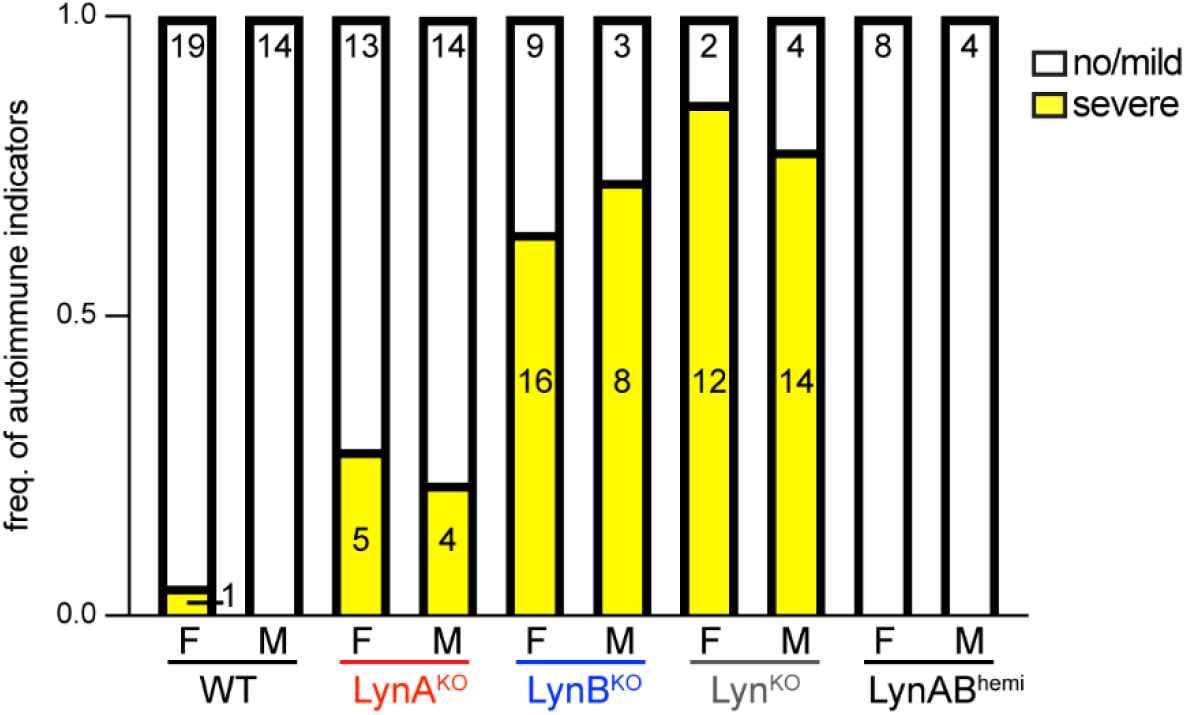
LynB^KO^ increases the incidence of autoimmune disease in male and female mice. Frequency of autoimmune indicators in male and female mice aged 8 months. Numbers represent individual tests, which may result in multiple counts from an individual. There were no significant differences between males and females within any genotype, as assessed via two-sided Fisher’s exact test.

